# Egg-stage desiccation reduces developmental recovery and reveals strain-dependent *Wolbachia*-associated costs in the Mediterranean fruit fly, *Ceratitis capitata*

**DOI:** 10.64898/2026.04.21.719827

**Authors:** M Kamilari, G Giannatos, G Tsiamis, AA Augustinos

## Abstract

The Mediterranean fruit fly (medfly) (*Ceratitis capitata* (Wiedemann, 1824) is a major agricultural pest, and egg desiccation is a critical constraint during handling and mass-rearing, as even short periods without moisture may compromise developmental success and downstream adult performance. The *Wolbachia*–medfly symbiosis is a relatively recently established artificial association, generated less than three decades ago using *Rhagoletis cerasi* as the *Wolbachia* donor. In this study, we evaluated the effects of egg-stage desiccation on developmental success and subsequent adult performance in three medfly lines differing in *Wolbachia* status: the uninfected Benakeion line, the *w*Cer2-infected 88.6 line, and the *w*Cer4-infected S10.3 line. Eggs were exposed to desiccation for 0–24 h at 4-h intervals before transfer to larval diet, and hatching, pupation, and adult emergence were recorded. We additionally assessed adult survival under stress for flies emerging from the 0, 8, and 10 h egg-desiccation treatments. Under control conditions, Benakeion showed the highest hatching and developmental recovery, S10.3 the lowest, and 88.6 intermediate performance. Across all strains, short desiccation exposures were comparatively well tolerated, whereas prolonged exposure sharply reduced hatching, pupation, and adult emergence, with the clearest decline at 20–24 h. Strain-dependent differences were expressed mainly at the hatching stage, while later developmental transitions were more similar among strains once larvae had hatched. In the adult follow-up, strain, rather than moderate egg-stage desiccation, was the main determinant of short-term survival and survival under extreme stress, with S10.3 again showing the weakest performance. These results indicate that *Wolbachia*-associated fitness costs in medfly are strain dependent and that egg-stage desiccation primarily acts at the embryonic bottleneck. Beyond providing insight into the *Wolbachia*–medfly artificial symbiosis, our findings are directly relevant to egg-handling and strain-evaluation protocols in medfly mass-rearing systems.

## 1. Introduction

Water balance is a central determinant of insect performance, affecting survival, development, behaviour, and ecological persistence. In terrestrial insects, desiccation stress arises when evaporative water loss exceeds the capacity for water acquisition and physiological compensation, and coping with this imbalance depends on integrated responses involving the cuticle, excretory system, gut, metabolism, and behaviour. Because all terrestrial insects experience fluctuating temperature and humidity, they may encounter desiccation stress continuously or intermittently, with consequences ranging from reduced activity and altered behaviour to developmental disruption and mortality (Crowe, 2015; McCluney, 2017; McCluney et al., 2012; Moss et al., 2024; Weldon et al., 2018, 2016). At the physiological level, water loss can lead to a hyperosmotic state that impairs protein folding, enzyme activity, and cellular homeostasis, making osmoregulation and dehydration tolerance fundamental components of insect resilience (Addo-Bediako et al., 2001; Chown et al., 2011; Halberg and Denholm, 2024; Thorat and Nath, 2018; Weldon et al., 2019, 2018). Although the general principles of insect osmoregulation are well established, responses to desiccation remain insufficiently characterized for many economically important non-model species.

Insects exhibit diverse physiological and behavioural strategies for coping with water scarcity. At the physiological level, variation in cuticular permeability, renal and digestive function, and metabolic responses can all contribute to the maintenance of water and ion balance under dehydration stress (Chown et al., 2011; Halberg and Denholm, 2024; Lee, 2002; Thorat and Nath, 2018). At the behavioural level, insects may modify activity patterns, seek out more humid microhabitats, or alter reproductive investment in response to desiccating conditions(Bartlett et al., 2019; Bujan et al., 2016; Johnson et al., 2023). Such strategies are ecologically significant because temperature and humidity often fluctuate together in nature, and periods of high temperature coupled with low relative humidity can reduce survival and population persistence if water loss cannot be compensated (Fischer and Kirste, 2018; Holmes and Benoit, 2019; Schmidt et al., 2018; Simmons et al., 2023; Wang et al., 2021). Understanding how insects respond to desiccation stress is therefore important not only from a physiological perspective, but also for interpreting species distributions, host use, and performance in changing environments.

The Mediterranean fruit fly (medfly), *Ceratitis capitata* (Wiedemann), is among the most economically important tephritid pests worldwide and a major target of area-wide control programmes. Native to temperate regions but now widely established in subtropical and tropical areas, medfly continues to expand its geographic range, with its seasonal biology and invasion success shaped by temperature, humidity, and host availability (Szyniszewska *et al*, 2024; Papadogiorgou *et al*, 2024; Szyniszewska & Tatem, 2014). Because the species exploits fleshy fruit and other humid microhabitats for oviposition and larval development, the egg and larval stages are especially sensitive to water balance, whereas adults may experience water stress outside the host or under suboptimal environmental conditions (Eskafi and Fernandez, 1990; Giunti et al., 2023; Sinclair et al., 2024; Weldon et al., 2016). Earlier work showed that medfly eggs are highly sensitive to low-humidity exposure, with hatchability declining sharply under severe drying conditions, indicating that embryonic development constitutes a major bottleneck in the response to water loss (Shoukry and Hafez, 1979). From an applied perspective, egg survival and the quality of resulting adults are directly relevant to the efficiency of mass-rearing systems and to standard quality-control frameworks used in tephritid production programmes (FAO/IAEA/USDA., 2019).

Medfly is also a tractable and highly informative model within Tephritidae. Its economic importance and cosmopolitan distribution have made it a key system in genetics, molecular biology, biotechnology, behaviour, and environmentally friendly insect control approaches (Bravo and Zucoloto, 1998; Dyck et al., 2021; Franz et al., 2021; Giunti et al., 2023). Medfly was the first fruit fly in which polytene chromosome maps were developed, providing evidence for the conservation of Muller elements in Diptera and facilitating later cytogenetic and genomic advances (Zacharopoulou et al., 2017). It was also the first non-drosophilid insect to be genetically transformed (Loukeris et al., 1995; Zwiebel et al., 1995). In addition, medfly has served as a model species for the development of the sterile insect technique (SIT), including extensive work on artificial rearing and genetic sexing strains for male-only releases (Franz et al., 2021). These characteristics make medfly particularly suitable for examining how physiological stress at the egg stage may affect developmental success and the biological quality of resulting adults.

The medfly also provides a useful system for studying the fitness consequences of artificial *Wolbachia* associations. *Wolbachia* is a widespread intracellular α-proteobacterium associated with diverse reproductive phenotypes in arthropods, including parthenogenesis, feminization, male killing, and cytoplasmic incompatibility (CI) (Hilgenboecker et al., 2008; O’Neill and Karr, 1990; Saridaki and Bourtzis, 2010; Yen and Barr, 1971; Zug and Hammerstein, 2012). Stable transinfections with the *Wolbachia* strains *w*Cer2 and *w*Cer4 were established previously in *C. capitata*, using *Rhagoletis cerasi* as donor, in the context of exploring the possible exploitation of Wolbachia-induced CI in agricultural pest control. Both strains induce strong CI in the novel host, including unidirectional incompatibility against uninfected females and bidirectional incompatibility between the two infected lines (Zabalou et al., 2009, 2004). More broadly, medfly has become one of the best tephritid systems for evaluating both the biological and applied consequences of *Wolbachia* infection in fruit flies (Mateos et al., 2020).

However, the medfly–Wolbachia symbiosis does not come without fitness costs. Initial and follow-up studies have shown that the effects of *Wolbachia* on medfly biology are strain dependent, with infected lines showing reduced hatching, altered immature performance, increased sensitivity to elevated temperatures, and reduced male sexual signalling, among other effects (Dionysopoulou et al., 2020; Kyritsis et al., 2022, 2019; Sarakatsanou et al., 2011; Zabalou et al., 2009, 2004). In particular, the *w*Cer4-infected line S.10.3 has repeatedly shown stronger costs than the *w*Cer2-infected line 88.6, indicating that host genotype and *Wolbachia* strain interact to shape life-history traits and tolerance to challenging developmental conditions (Dionysopoulou et al., 2020; Kyritsis et al., 2022, 2019). Because transient drying of eggs may occur during handling and transfer in rearing facilities, any Wolbachia-associated reduction in desiccation tolerance could influence both developmental recovery and the biological quality of resulting adults. Despite this, the interaction between egg-stage desiccation and Wolbachia infection in medfly has not been examined directly.

In the present study, we investigated whether desiccation duration affects hatching, pupation, and adult emergence in three medfly lines differing in Wolbachia status, namely the uninfected Benakeion line and the *w*Cer2- and *w*Cer4-infected lines 88.6 and S.10.3. We also examined whether moderate egg-stage desiccation has downstream effects on adult survival under stress. We predicted that prolonged egg desiccation would reduce developmental recovery, that the strongest strain-dependent divergence would be expressed at hatching, and that the *w*Cer4-infected line would show the greatest performance cost. By addressing these questions, the study links stress physiology with an applied problem in medfly strain evaluation and egg-handling protocols.

## 2. Materials and Methods

### 2.1 Insect strains and rearing conditions

All experiments were conducted at the Department of Plant Protection of Patras (ELGO-DIMITRA, Greece) using *C. capitata* strains maintained routinely in the insectary. Adults were kept in plastic-framed cages (30 × 30 × 10 cm) with mesh-covered sides for oviposition under controlled conditions of 25 ± 1°C, 55% relative humidity (RH), and a 14:10 h light:dark photoperiod.

Three medfly lines differing in Wolbachia status were used: (i) Benakeion, a long-established laboratory line free of Wolbachia; (ii) 88.6, a Benakeion-derived transinfected line carrying the *w*Cer2 Wolbachia strain; and (iii) S.10.3, a Benakeion-derived transinfected line carrying the *w*Cer4 Wolbachia strain. The transinfected lines were originally generated by artificial transfer of Wolbachia from *Rhagoletis* cer*asi* into the medfly background, as described previously (Zabalou et al., 2004). The infected lines have since been maintained through inbreeding under laboratory conditions.

### 2.2 Verification of Wolbachia infection status

Prior to the experiments, the Wolbachia status of all three strains was verified by diagnostic PCR. Genomic DNA was extracted from six adult individuals per strain using a CTAB-based protocol (Kamilari et al., 2025). DNA quality was first checked using universal 12S rRNA primers (Hanner and Fugate, 1997). Wolbachia infection status was then determined using strain-specific primer pairs targeting the *wsp* variants diagnostic for *w*Cer2 and *w*Cer4 (Arthofer *et al*, 2009). PCR products were separated on 2% agarose gels and visualized under UV illumination. All individuals yielded the expected amplification pattern, confirming the infection status of the three experimental lines.

### 2.3 Egg collection and desiccation treatments

Eggs were collected during the early morning (08:00–11:00 h) from cages at peak oviposition activity (days 4–7). Freshly collected eggs were transferred onto dry black filter paper and arranged in replicate batches of approximately 30 eggs each. For each strain, replicate egg batches were assigned to one of seven desiccation treatments: 0, 4, 8, 12, 16, 20, or 24 h of exposure. Finally, replicate batches from all runs were pooled for analysis. For the control treatment (0 h), three replicate egg batches per strain were transferred immediately to Petri dishes containing standard larval diet. For the desiccation treatments, eggs remained on dry filter paper without an external moisture source for the assigned duration and were then transferred to larval diet. Thus, for each strain, developmental performance was assessed after a graded series of egg-stage desiccation exposures from 0 to 24 h at 4-h intervals (**Fig.1).**

**Figure 1.**
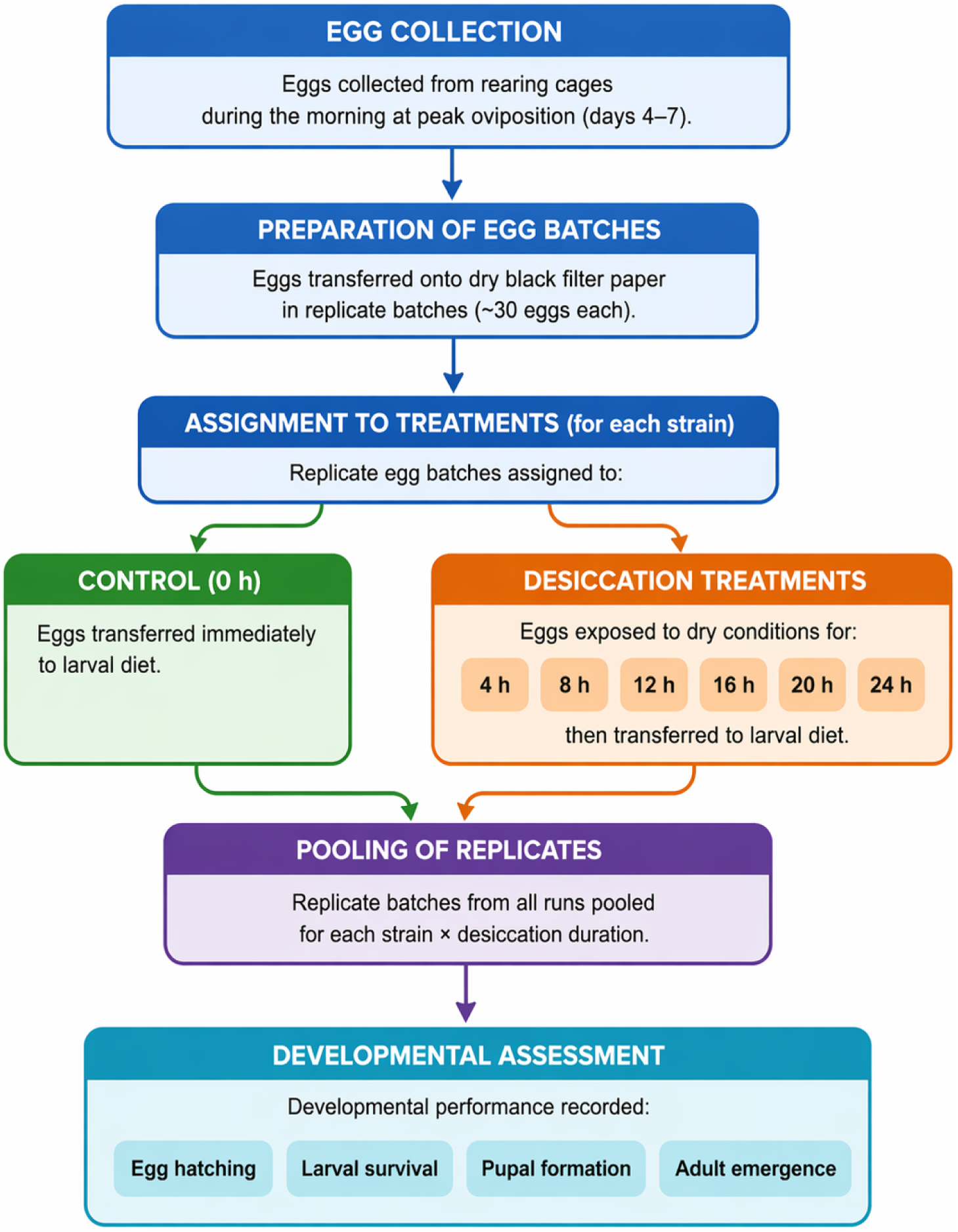
Experimental workflow for egg-stage desiccation treatments and developmental assessment in *Ceratitis capitata*.

### 2.4 Assessment of hatching, pupation, and adult emergence

Hatching was scored four days after oviposition. At that point, each Petri dish was transferred to an individual plastic box containing sand as pupation substrate. Seven days later, pupae were recovered by sieving, counted, and transferred to new Petri dishes for adult emergence. Adult emergence was considered complete after 11 days, and the total number of emerged adults was recorded.

For each replicate, developmental performance was quantified using both egg-based and conditional-stage metrics. The following response variables were calculated: i) Hatching (%) = number of hatched eggs / initial number of eggs × 100; ii) Pupation on eggs (%) = number of pupae / initial number of eggs × 100; iii) Emergence on eggs (%) = number of emerged adults / initial number of eggs × 100; iv) Pupation on hatched eggs (%) = number of pupae / number of hatched eggs × 100; v) Emergence on pupae (%) = number of emerged adults / number of pupae × 100. This two-level approach allowed us to distinguish overall developmental recovery from stage-specific survival after successful completion of the preceding stage.

### 2.5 Effects of egg-stage desiccation on adult survival under stress

To evaluate whether moderate egg-stage desiccation had downstream effects on adult quality, a targeted adult follow-up assay was conducted using flies emerging from three egg treatments: control (0 h), 8 h desiccation, and 10 h desiccation. The 8 h and 10 h treatments were selected intentionally based on the main developmental assay, which indicated that the transition from limited to more pronounced detrimental effects occurred within the 8–12 h interval. Thus, 8 h was included as the upper bound of relatively tolerated exposure, whereas 10 h was selected as an intermediate exposure within this biologically relevant transition zone. Longer desiccation exposures were not included in the adult assay because they yielded too few surviving individuals for a meaningful assessment of post-emergence performance.

Following emergence, 20–40 adult flies per strain and treatment group were randomly selected and individually housed in vials without access to food or water. Each fly’s survival under extreme stress, defined as the number of hours from adult emergence to death, was recorded. In addition, survival at 48 h post-emergence was scored as a binary variable (1 = alive at 48 h; 0 = dead before 48 h), as a practical proxy of adult biological quality in line with quality-control approaches used in tephritid mass-rearing programmes. Mortality was checked twice daily at 12 h intervals (FAO/IAEA/USDA., 2019).

### 2.6 Statistical analysis

As a preliminary step, we analyzed the three non-desiccated control strains to establish baseline developmental performance under unstressed conditions. For each strain, hatching, pupation, and adult emergence were calculated both relative to the initial number of eggs and relative to the preceding developmental stage.

To assess the effects of egg-stage desiccation on development, we analyzed hatching, pupation, and adult emergence across the full desiccation series (0–24 h) in the three medfly strains. The primary analysis focused on egg-based endpoints, namely hatching (%), pupation on eggs (%), and emergence on eggs (%), because these variables capture overall developmental recovery from the initial egg cohort. Conditional-stage endpoints, namely pupation on hatched eggs (%) and emergence on pupae (%), were analyzed separately to determine whether strain effects persisted after successful completion of earlier developmental transitions.

Normality of residuals was assessed using the Shapiro–Wilk test and homogeneity of variance using Levene’s test. When assumptions of normality and homoscedasticity were met, parametric analyses were used. Specifically, baseline control comparisons among strains were evaluated by one-way ANOVA, followed by Tukey’s honestly significant difference (HSD) test for pairwise contrasts. When assumptions were not met, non-parametric tests were applied. In those cases, differences among strains or among desiccation durations were tested using the Kruskal–Wallis test, followed by Dunn’s post hoc comparisons with Bonferroni correction. For the desiccation-duration analyses, hatching, pupation on eggs, and emergence on eggs were compared across the seven time points (0, 4, 8, 12, 16, 20, and 24 h). For strain-level comparisons across treatments, hatching, pupation on hatched eggs, and emergence on pupae were compared among Benakeion, 88.6, and S.10.3.

For the adult follow-up assay, two response variables were analyzed: survival at 48 h and total survival under extreme stress under stress. This targeted assay included only the control, 8 h, and 10 h egg-desiccation treatments, because these exposure levels were chosen to represent the biologically relevant transition zone identified by the main developmental dataset while still providing enough surviving adults for downstream analysis. Survival at 48 h was analyzed using binomial logistic regression, with strain, egg-desiccation treatment, and their interaction included as fixed effects. Adult survival under extreme stress was analyzed using a two-way ANOVA with strain and egg-desiccation treatment as factors after checking model assumptions with the Shapiro–Wilk and Levene’s tests. When ANOVA assumptions were not met, non-parametric alternatives (Kruskal–Wallis test followed by Dunn’s post hoc test with Bonferroni correction) were applied.

All statistical analyses were performed in SPSS v16. Dunn’s post hoc tests were carried out using StatsCalculators (2025) (“StatsCalculators.com - Free Online Statistics Calculators,” n.d.). The significance threshold was set at α = 0.05, with Bonferroni adjustment applied where appropriate for multiple pairwise comparisons.

## 3. Results

### 3.1. Wolbachia-status verification

All individuals screened from the three medfly strains yielded the expected amplicon pattern in the diagnostic PCR assays, confirming the Wolbachia status of the experimental lines: Benakeion as uninfected, 88.6 as infected with *w*Cer2, and S.10.3 as infected with *w*Cer4.

### 3.2. Response to egg-stage desiccation stress

#### 3.2.1. Baseline developmental performance of the three strains under control conditions

Under non-desiccating conditions, the three *Ceratitis capitata* strains differed markedly in overall developmental performance (**Table 1**; **Figure 2**). Benakeion showed the highest hatching, pupation, and adult emergence when these endpoints were calculated relative to the initial number of eggs, whereas S.10.3 consistently showed the lowest values and 88.6 was intermediate. Mean hatching reached 94.47 ± 3.87% in Benakeion, compared with 73.37 ± 1.82% in 88.6 and 58.33 ± 3.82% in S.10.3. A similar ranking was observed for pupation on eggs and emergence on eggs.

**Figure 2.**
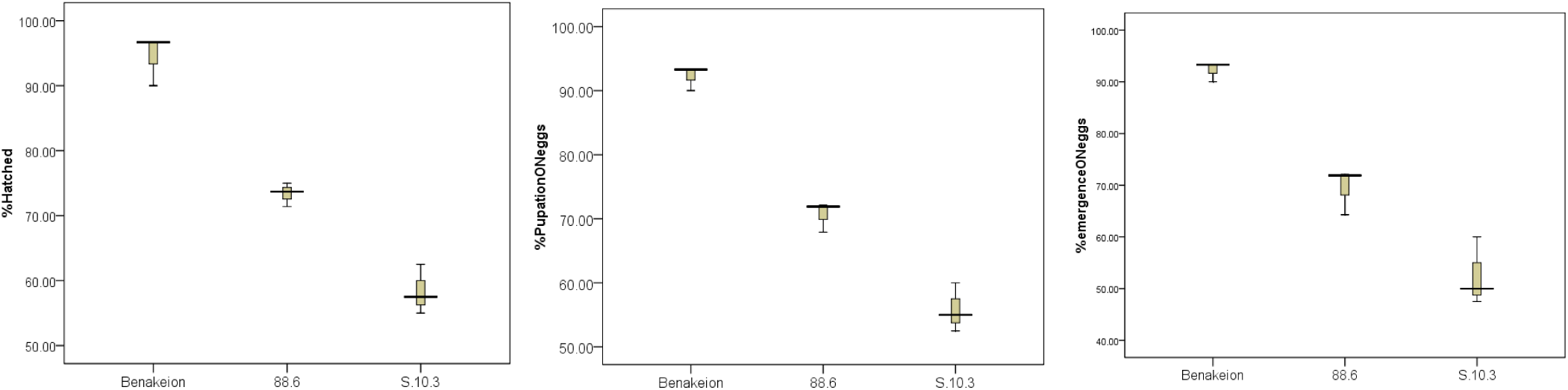
Baseline developmental performance under control conditions. Panels show mean percentage hatching, pupation, and adult emergence in the three strains under non-desiccating conditions; the clearest between-strain separation is observed for egg-based endpoints.

**Table 1.** Baseline control performance summary (mean ± SD) for the three Ceratitis capitata strains. Mean ± SD values for hatching, pupation, and adult emergence in the Benakeion, 88.6 (wCer2), and S.10.3 (wCer4) strains under non-desiccating conditions. Developmental success is expressed both relative to the initial number of eggs and relative to the preceding developmental stage, allowing comparison of overall developmental recovery and conditional-stage survival.

| Strain | Hatching (%) | Pupation on eggs (%) | Emergence on eggs (%) | Pupation on hatched eggs (%) | Emergence on pupae (%) |
| --- | --- | --- | --- | --- | --- |
| Benakeion | 94.47 ± 3.87 | 92.20 ± 1.91 | 92.20 ± 1.91 | 97.73 ± 1.96 | 100.00 ± 0.00 |
| 88.6 | 73.37 ± 1.82 | 70.63 ± 2.37 | 69.43 ± 4.45 | 96.23 ± 1.31 | 98.23 ± 3.06 |
| S.10.3 | 58.33 ± 3.82 | 55.83 ± 3.82 | 52.50 ± 6.61 | 95.77 ± 4.35 | 93.87 ± 6.90 |

When developmental success was expressed relative to the preceding developmental stage, differences among strains were greatly reduced. Pupation among hatched larvae remained high in all strains, ranging from 95.77 ± 4.35% in S.10.3 to 97.73 ± 1.96% in Benakeion, while emergence among pupae ranged from 93.87 ± 6.90% in S.10.3 to 100.00 ± 0.00% in Benakeion (**Table 1**).

The inferential analyses supported the same pattern (**Table 2**). Significant strain differences were detected for hatching, pupation on eggs, and emergence on eggs, whereas no significant strain effect was detected for pupation on hatched eggs or emergence on pupae. Thus, under control conditions, the major among-strain divergence was concentrated at the egg-to-larva transition rather than in later developmental stages. Full descriptive statistics are provided in **Table S1**, and the corresponding conditional-stage control profiles are shown in **Figure S1**.

**Table 2.** Summary of baseline control-condition comparisons among the three Ceratitis capitata strains. Statistical comparison of developmental endpoints under control conditions. The table summarizes the test used, test statistic, p-value, and overall outcome for hatching, pupation, and adult emergence expressed either relative to the initial number of eggs or to the preceding developmental stage.

| Endpoint | Test | Statistic | p-value | Interpretation |
| --- | --- | --- | --- | --- |
| Hatching (%) | One-way ANOVA | F = 90.211 | <0.001 | All three strains differ; Benakeion > 88.6 > S.10.3. |
| Pupation on hatched eggs (%) | One-way ANOVA | F = 0.388 | 0.695 | No significant strain effect. |
| Pupation on eggs (%) | One-way ANOVA | F = 126.331 | <0.001 | All three strains differ. |
| Emergence on pupae (%) | One-way ANOVA | F = 1.576 | 0.282 | No significant strain effect. |
| Emergence on eggs (%) | One-way ANOVA | F = 53.188 | <0.001 | All three strains differ. |

#### 3.2.2. Effects of desiccation duration on developmental success calculated against the initial egg number

Desiccation duration had a strong negative effect on developmental success across all three strains. When hatching, pupation, and adult emergence were assessed relative to the initial number of eggs, Kruskal-Wallis tests showed highly significant effects of desiccation duration for all three endpoints (**Table 3**; **Figure 3**).

**Figure 3.**
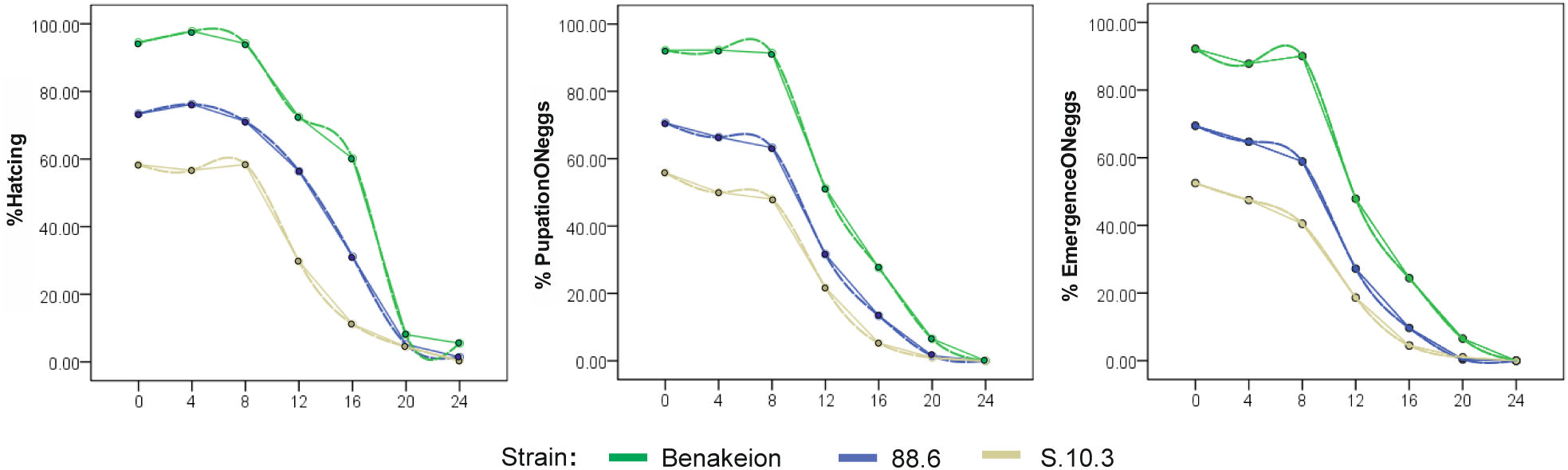
Estimated marginal means or corresponding summary profiles for hatching, pupation on eggs, and adult emergence on eggs across the seven egg-desiccation treatments (0, 4, 8, 12, 16, 20, and 24 h). The figure depicts the overall temporal pattern of developmental recovery across the desiccation gradient, highlighting the relatively limited effects of short exposures and the marked decline associated with prolonged desiccation.

**Table 3.** Summary of desiccation-duration effects on egg-based developmental endpoints in Ceratitis capitata. Kruskal–Wallis results for hatching, pupation on eggs, and adult emergence on eggs across the seven egg-desiccation treatments (0, 4, 8, 12, 16, 20, and 24 h). The table reports the test statistics, degrees of freedom, p-value, and the principal post hoc conclusion for each endpoint.

| Endpoint | Kruskal–Wallis $\chi^2$ | df | p-value | Key post hoc conclusion |
| --- | --- | --- | --- | --- |
| Hatching (%) | 47.098 | 6 | <0.001 | No difference among 0, 4, and 8 h; strong decline at 20 and 24 h relative to early time points. |
| Pupation on eggs (%) | 53.506 | 6 | <0.001 | Short exposures are tolerated; prolonged desiccation sharply reduces success. |
| Emergence on eggs (%) | 53.074 | 6 | <0.001 | Long exposures, especially 20–24 h, cause strong loss of developmental recovery. |

Across strains, developmental performance remained comparatively stable during short desiccation exposures and then declined progressively with increasing duration. No significant differences were detected among the 0, 4, and 8 h treatments for hatching, pupation on eggs, or emergence on eggs. By contrast, prolonged exposure resulted in a marked loss of developmental recovery, with the clearest and most consistently supported reductions observed at 20 and 24 h. Thus, medfly eggs tolerated short desiccation exposures relatively well under the conditions tested, whereas extended water deprivation sharply reduced developmental success. Full Dunn pairwise comparisons for the desiccation-duration analysis are provided in **Table S2**.

#### 3.2.3. Strain-specific differences under desiccation and stage-specific conditional survival

Across all desiccation treatments combined, strain significantly affected hatching success, whereas effects on downstream developmental transitions were weaker. For hatching percentage, the Kruskal–Wallis test was significant and Dunn’s post hoc comparisons showed that Benakeion differed significantly from S.10.3, while 88.6 did not differ significantly from either strain. No significant strain differences were detected for pupation among hatched larvae or emergence among pupae.

Benakeion consistently showed the highest developmental recovery across desiccation treatments, S.10.3 the lowest, and 88.6 intermediate values. However, once larvae successfully hatched, subsequent completion of pupation and adult emergence was broadly similar among strains. These findings indicate that the strain-dependent component of the desiccation response was expressed primarily at the level of egg viability and hatching. The full sequential-stage statistical results are provided in **Table S3**, and the corresponding conditional-stage profiles across desiccation treatments are shown in **Figure S3**.

#### 3.2.4. Adult survival under extreme stress (no food and water provision) following egg-stage desiccation

Because the main developmental assay indicated that the biologically relevant transition from limited to more pronounced detrimental effects occurred between 8 and 12 h of egg desiccation, the adult follow-up focused on control, 8 h, and an intermediate 10 h exposure. This design allowed evaluation of post-emergence performance within the most informative applied range while avoiding longer egg-desiccation treatments that yielded too few surviving adults for meaningful downstream analysis.

To assess whether egg-stage desiccation had downstream effects on adult quality, we evaluated survival at 48 h after emergence and total adult survival under extreme stress under starvation and dehydration. Survival at 48 h remained high in Benakeion and 88.6 across treatments, whereas S.10.3 showed lower survival, particularly after egg desiccation **(Table 4**). Logistic regression identified a significant overall effect of strain on 48 h survival. Relative to S.10.3, the odds of survival were significantly higher in 88.6 and even higher in Benakeion (Table 5). Full logistic regression coefficients and confidence intervals are provided in **Table S4.**

**Table 4.** Adult descriptive outcomes after moderate egg-stage desiccation in three Ceratitis capitata strains. Percentage of adults alive at 48 h, mean adult survival under extreme stress, and median adult survival under extreme stress for the Benakeion, 88.6, and S.10.3 strains after egg treatments of 0, 8, and 10 h desiccation. Adults were maintained without food or water after emergence to assess post-emergence performance under stress.

| Strain | Treatment | Alive at 48 h (%) | Mean survival under extreme stress (h) | Median survival under extreme stress (h) |
| --- | --- | --- | --- | --- |
| Benakeion | Control | 95 | 70.86 | 72 |
| Benakeion | 8 h desiccation | 93 | 70.57 | 72 |
| Benakeion | 10 h desiccation | 95 | 73.43 | 72 |
| 88.6 | Control | 95 | 66.00 | 66 |
| 88.6 | 8 h desiccation | 100 | 73.80 | 72 |
| 88.6 | 10 h desiccation | 90 | 72.00 | 72 |
| S.10.3 | Control | 91 | 66.00 | 66 |
| S.10.3 | 8 h desiccation | 73 | 63.82 | 60 |
| S.10.3 | 10 h desiccation | 82 | 61.64 | 60 |

**Table 5:** Statistical summary of the adult follow-up assays for 48 h survival and survival under extreme stress after egg-stage desiccation. Summary of the statistical models used to assess post-emergence performance in adults originating from eggs exposed to 0, 8, or 10 h desiccation. The table reports the model used, the principal test results, and the overall outcome for survival at 48 h and total adult survival under extreme stress under stress.

| Outcome | Model | Key result | Interpretation |
| --- | --- | --- | --- |
| 48 h survival | Logistic regression | Omnibus $\chi^2 = 12.52$, $p = 0.002$; OR 88.6 vs S.10.3 = 3.59 ($p = 0.015$); OR Benakeion vs S.10.3 = 9.00 ($p = 0.005$). | Short-term adult survival differs mainly by strain, with S.10.3 performing worst. |
| 48 h survival | GLM terms | Treatment $p = 0.625$; strain × treatment $p = 0.944$. | No significant treatment effect or strain × treatment interaction was detected in the current model output. |
| survival under extreme stress | Two-way ANOVA | Strain: $F(2,244) = 13.82$, $p < 0.001$; treatment: $F(2,244) = 2.04$, $p = 0.132$. | Strong strain effect; no clear downstream treatment effect. |

Adult survival under extreme stress showed a similar pattern (**Figure 4**). Mean survival under extreme stress remained relatively stable across treatments within Benakeion and 88.6, whereas S.10.3 consistently showed lower values. The statistical analyses supported a significant main effect of strain on adult survival under extreme stress, whereas the main effect of egg-desiccation treatment was not significant (**Table 5**). Likewise, the currently reported model output for 48 h survival did not support a significant treatment effect or a significant strain × treatment interaction.

**Figure 4.**
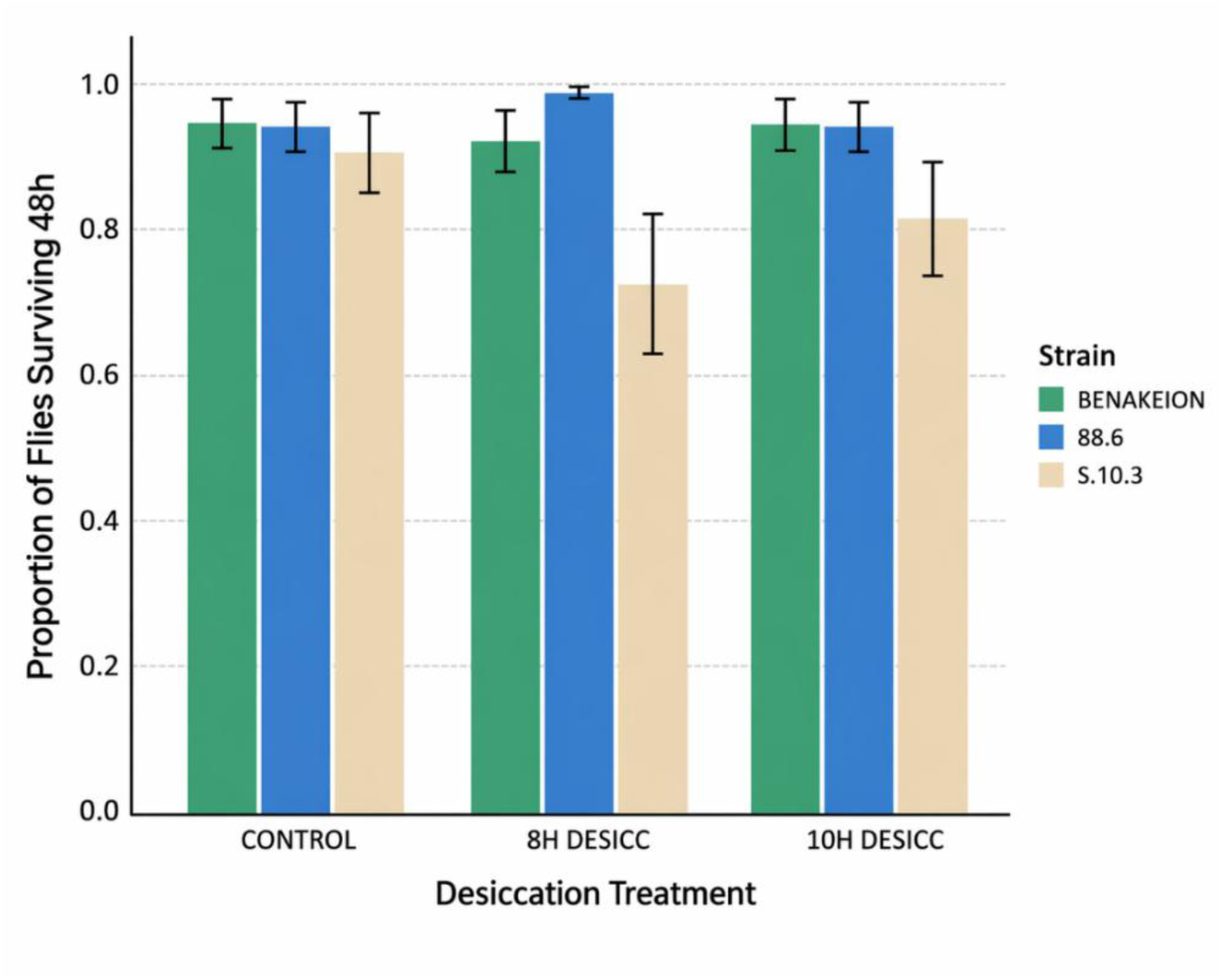
Mean adult survival under extreme stress by strain and egg-desiccation treatment. Benakeion and 88.6 maintain comparatively robust adult performance across treatments, whereas S.10.3 shows consistently lower survival under extreme stress.

Taken together, the adult follow-up assays indicate that post-emergence performance under stress was determined mainly by strain identity rather than by moderate egg-stage desiccation. Among the three lines, S.10.3 consistently showed the weakest adult performance, whereas Benakeion and 88.6 were comparatively robust.

## 4. Discussion

Egg-stage desiccation had a strong negative effect on developmental recovery in *Ceratitis capitata*, but this effect was not uniform across strains. Across the three lines examined here, developmental performance remained comparatively stable during short desiccation exposures and then declined sharply after prolonged drying, with the clearest reductions observed at the longest exposure times. At the same time, the three strains differed substantially in both baseline and stress-associated performance: Benakeion consistently showed the highest values, S.10.3 the lowest, and 88.6 an intermediate phenotype. Importantly, the main among-strain divergence was concentrated at the hatching stage. Once larvae had hatched successfully, subsequent pupation and adult emergence were much more similar among strains. This indicates that egg viability and successful completion of embryogenesis represent the major bottleneck in the response to desiccation in this system (Halberg and Denholm, 2024; Shoukry and Hafez, 1979).

This interpretation is consistent with general principles of insect water balance. Desiccation stress arises when water loss exceeds the insect’s capacity for physiological compensation, and the ability to withstand such stress depends on the coordinated function of the cuticle, excretory system, gut, and metabolic homeostasis. In egg stages, where developmental progression depends on maintaining sufficient hydration for embryogenesis, water loss is expected to impose particularly strong constraints. Our data fit this framework well, because the clearest effect of desiccation was expressed before or at hatching rather than during later immature development. The results therefore support the view that, under the conditions tested here, egg-stage dehydration primarily compromises embryonic viability rather than causing a generalized reduction in survival across all later developmental stages (Halberg and Denholm, 2024).

The temporal pattern of the response is also biologically informative. Short desiccation exposures caused little measurable reduction in hatching, pupation, or adult emergence, whereas prolonged exposure led to a marked decline in developmental recovery. This pattern suggests a threshold-like deterioration in performance rather than a strictly linear response across time. Earlier work on medfly biology reported that egg desiccation under severe low-humidity conditions causes a strong loss of hatchability, with little or no recovery after prolonged exposure (Chapman et al., 1998; McDermott and Mullens, 2014). Although the exposure conditions in the present study were less extreme than those historical assays, the qualitative pattern is consistent: medfly eggs tolerate only a limited period of drying before developmental recovery collapses. The present results therefore extend earlier observations by showing that this sensitivity is also modulated by *Wolbachia* strain background in transinfected medfly lines.

The baseline control data reinforce this interpretation. Even in the absence of desiccation, Benakeion outperformed both *Wolbachia*-infected lines when developmental success was calculated from the initial number of eggs, whereas differences among strains largely disappeared when pupation and emergence were expressed relative to the preceding developmental stage. This pattern suggests that the main cost associated with the infected lines, particularly S.10.3, is not a generalized reduction in performance across all immature stages but rather a stronger effect on the earliest transition from egg to larva. Among the two infected lines, S.10.3, which carries *w*Cer4, consistently showed the weakest performance, with the lowest baseline hatching, the lowest developmental recovery under desiccation, and the poorest adult survival and survival under extreme stress under stress. In contrast, the 88.6 line carrying *w*Cer2 was much closer to the uninfected Benakeion line, especially in the adult assays. The most parsimonious interpretation is therefore that the cost of the artificial medfly–Wolbachia association is strain dependent, with *w*Cer4 imposing a stronger burden on the host than *w*Cer2 under the conditions tested here. The present data do not resolve mechanism, but they are compatible with the hypothesis that Wolbachia strain identity affects embryonic robustness and thereby alters the capacity of eggs to withstand temporary water loss. This interpretation is consistent with the original medfly transinfection work, which demonstrated stable establishment and strong cytoplasmic incompatibility in both lines, and with later studies showing that the phenotypic consequences of infection differ substantially between *w*Cer2 and *w*Cer4 backgrounds (Kyritsis et al., 2022, 2019; Zabalou et al., 2004).

From an applied perspective, this result is relevant beyond the specific question of Wolbachia biology. In medfly rearing systems, the egg stage is operationally important because egg collection, temporary holding, transfer, and handling conditions can influence downstream productivity. The present findings indicate that short periods of drying may be tolerated under the laboratory conditions used here, whereas prolonged desiccation rapidly reduces developmental yield. They also show that host genotype and symbiotic background can influence this response. This is important because mass-rearing systems, and especially programmes that evaluate novel lines for operational use, benefit from knowing whether hidden fitness costs are expressed at the egg stage. The study therefore contributes directly to the biological basis for egg-handling and strain-evaluation protocols in medfly rearing facilities (Bourtzis and Vreysen, 2021; Elaini et al., 2020; FAO/IAEA/USDA., 2019).

The adult stress assay was designed as a targeted follow-up focused on the most informative applied exposure range, adding an important second layer to the interpretation. Specifically, adults were assessed after control, 8 h, and 10 h egg desiccation because the main developmental dataset indicated that the 8–12 h interval represented the transition from relatively tolerated exposure to increasingly detrimental effects on developmental recovery. This is also the exposure range of greatest practical interest for facilities, because it captures the boundary at which temporary egg handling may begin to compromise developmental yield while still permitting enough recovery to evaluate the biological quality of resulting adults. Selecting an intermediate 10 h treatment therefore allowed assessment of downstream adult performance within this biologically relevant window, while avoiding longer desiccation durations that yielded too few survivors for informative adult-quality analysis. Within this range, adults emerging from desiccated eggs did not show a clear reduction in 48 h survival or total survival under extreme stress attributable to treatment itself. Instead, post-emergence performance was explained mainly by strain identity, with S.10.3 again showing the lowest robustness and Benakeion and 88.6 performing better. This suggests that, within the range of egg desiccation examined in the adult follow-up, survivors did not carry a strong detectable downstream penalty in this particular assay. That point is important because survival under stress is commonly used as a practical biological-quality indicator in tephritid rearing, including in the evaluation of sterile males destined for SIT release purposes. Thus, the present results suggest that moderate egg-stage desiccation reduced developmental yield, but did not produce a strong additional deterioration in this adult-quality proxy among those individuals that completed development (FAO/IAEA/USDA., 2019; Shelly, 2018).

With respect to IIT-based or broader Wolbachia-based control strategies, the present study is best viewed as a contribution to understanding the biology and hidden costs of an artificially established medfly–Wolbachia association and to the evaluation of rearing-relevant traits before operational deployment. The data indicate that not all Wolbachia infections are equivalent from a host-fitness perspective and that *w*Cer2 may be less costly than *w*Cer4 under desiccation-related stress. However, whether such differences translate into operational advantages in control programmes will depend on programme-specific needs, the applicable regulatory framework, and biological performance under production and field conditions. For that reason, the emphasis of the present study should remain on artificial symbiosis and strain evaluation rather than on strong claims about immediate operational application (Bourtzis and Vreysen, 2021; Mateos et al., 2020; Saridaki and Bourtzis, 2010; Zabalou et al., 2009).

In summary, this study shows that egg-stage desiccation sharply reduces developmental recovery in *C. capitata* after prolonged exposure, that strain-specific differences are expressed mainly at the hatching stage, and that the *w*Cer4-infected line S.10.3 bears the strongest cost under both control and stress conditions. By contrast, the 88.6 line remains much closer to the uninfected background, particularly in adult performance. These findings refine our understanding of the fitness consequences of artificial Wolbachia associations in medfly and provide directly relevant information for the design of egg collection, temporary holding, and strain-evaluation protocols in mass-rearing systems (Cáceres et al., 2007; FAO/IAEA/USDA., 2019).

## Supporting information

Suuplementary Files

## Acknowledgements

The research project was supported by the Hellenic Foundation for Research and Innovation (H.F.R.I.) under the “3rd Call for H.F.R.I. Research Projects to support Post-Doctoral Researchers” (Project No: 7615_DESICCOME: *Desiccation Tolerance in metazoans using comparative transcriptomics*) & the European Union’s HORIZON research and Innovation Programme (Grant Aggr. 101059623_REACT: *Rapid elimination of invasive insect agricultural pest outbreaks by tackling them with sterile Insect Technique programs;* www.react-insect.eu).

## References

Addo-Bediako, A., Chown, S.L., Gaston, K.J., 2001. Revisiting water loss in insects: a large scale view. Journal of Insect Physiology 47, 1377–1388. 10.1016/S0022-1910(01)00128-7

Arthofer, W., Riegler, M., Schneider, D., Krammer, M., Miller, W.J., Stauffer, C., 2009. Hidden Wolbachia diversity in field populations of the European cherry fruit fly, Rhagoletis cerasi (Diptera, Tephritidae). Molecular Ecology 18, 3816–3830. 10.1111/j.1365-294X.2009.04321.x

Bartlett, J., Convey, P., Hayward, S.A.L., 2019. Not so free range? Oviposition microhabitat and egg clustering affects Eretmoptera murphyi (Diptera: Chironomidae) reproductive success. Polar Biol 42, 271–284. 10.1007/s00300-018-2420-4

Bourtzis, K., Vreysen, M.J.B., 2021. Sterile Insect Technique (SIT) and Its Applications. Insects 12. 10.3390/insects12070638

Bravo, I.S.J., Zucoloto, F.S., 1998. Performance and feeding behavior of Ceratitis capitata: comparison of a wild population and a laboratory population. Entomologia Experimentalis et Applicata 87, 67–72. 10.1046/j.1570-7458.1998.00305.x

Bujan, J., Yanoviak, S.P., Kaspari, M., 2016. Desiccation resistance in tropical insects: causes and mechanisms underlying variability in a Panama ant community. Ecology and Evolution 6, 6282–6291. 10.1002/ece3.2355

CABI Compendium Invasive Species [WWW Document], 2024. CABI Digital Library. URL https://www.cabidigitallibrary.org/product/QI (accessed 1.9.25).

Cáceres, C., Ramírez, E., Wornoayporn, V., Islam, S.M., Ahmad, S., 2007. a protocol for storage and long-distance shipment of mediterranean fruit fly (diptera: tephritidae) eggs. i. effect of temperature, embryo age, and storage time on survival and quality. flen 90, 103–109. 10.1653/0015-4040(2007)90%255B103:APFSAL%255D2.0.CO;2

Chapman, T., Miyatake, T., Smith, H.K., Partridge, L., 1998. Interactions of mating, egg production and death rates in females of the Mediterranean fruit fly, Ceratitis capitata. Proc Biol Sci 265, 1879–1894. 10.1098/rspb.1998.0516

Chown, S.L., Sørensen, J.G., Terblanche, J.S., 2011. Water loss in insects: an environmental change perspective. J Insect Physiol 57, 1070–1084. 10.1016/j.jinsphys.2011.05.004

Crowe, J.H., 2015. Anhydrobiosis: An Unsolved Problem with Applications in Human Welfare. Subcell Biochem 71, 263–280. 10.1007/978-3-319-19060-0_11

Dionysopoulou, N.K., Papanastasiou, S.A., Kyritsis, G.A., Papadopoulos, N.T., 2020. Effect of host fruit, temperature and Wolbachia infection on survival and development of Ceratitis capitata immature stages. PLoS One 15, e0229727. 10.1371/journal.pone.0229727

Dyck, V.A., Hendrichs, J., Robinson, A.S. (Eds.), 2021. Sterile Insect Technique: Principles and Practice in Area-Wide Integrated Pest Management, 2nd ed. CRC Press, Taylor and Francis Group, LLC, Boca Raton, FL.

Elaini, R., Asadi, R., Naish, N., Koukidou, M., Ahmed, M., 2020. Evaluation of Rearing Parameters of a Self-Limiting Strain of the Mediterranean Fruit Fly, Ceratitis capitata (Diptera: Tephritidae). Insects 11. 10.3390/insects11100663

Eskafi, F.M., Fernandez, A., 1990. Larval–Pupal Mortality of Mediterranean Fruit Fly (Diptera: Tephritidae) from Interaction of Soil, Moisture, and Temperature. Environmental Entomology 19, 1666–1670. 10.1093/ee/19.6.1666

FAO/IAEA/USDA., 2019. Product Quality Control for Sterile Mass-Reared and Released Tephritid Fruit Flies, Version 7.0. [WWW Document]. URL http://www.iaea.org/resources/manual/product-quality-control-for-sterile-mass-reared-and-released-tephritid-fruit-flies-version-70 (accessed 4.15.26).

Fischer, K., Kirste, M., 2018. Temperature and humidity acclimation increase desiccation resistance in the butterfly icyclus anynana. Entomologia Experimentalis et Applicata 166, 289–297. 10.1111/eea.12662

Franz, G., Bourtzis, K., Caceres, C., 2021. Practical and Operational Genetic Sexing Systems Based on Classical Genetic Approaches in Fruit Flies, an Example for other Species Amenable to Large-Scale Rearing for the Sterile Insect Technique, in: Dyck, V.A., Robinson, A.S., Hendrichs, J. (Eds.), Sterile Insect Technique. Principles and Practice in Area-Wide Integrated Pest Management. Springer, Dordrecht, The Netherlands, pp. 575–604.

Giunti, G., Benelli, G., Campolo, O., Canale, A., Kapranas, A., Liedo, P., De Meyer, M., Nestel, D., Ruiu, L., Scolari, F., Wang, X., Papadopoulos, N.T., 2023. Biology, ecology and invasiveness of the Mediterranean fruit fly, Ceratitis capitata: a review. Entomologia Generalis 1221–1239. 10.1127/entomologia/2023/2135

Halberg, K.V., Denholm, B., 2024. Mechanisms of Systemic Osmoregulation in Insects. Annual Review of Entomology 69, 415–438. 10.1146/annurev-ento-040323-021222

Hanner, R., Fugate, M., 1997. Branchiopod Phylogenetic Reconstruction from 12s Rdna Sequence Data. Journal of Crustacean Biology 17, 174–183. 10.1163/193724097X00188

Hilgenboecker, K., Hammerstein, P., Schlattmann, P., Telschow, A., Werren, J.H., 2008. How many species are infected with Wolbachia? – a statistical analysis of current data. FEMS Microbiol Lett 281, 215–220. 10.1111/j.1574-6968.2008.01110.x

Holmes, C.J., Benoit, J.B., 2019. Biological Adaptations Associated with Dehydration in Mosquitoes. Insects 10, 375. 10.3390/insects10110375

Johnson, M.G., Alvarez, K., Harrison, J.F., 2023. Water loss, not overheating, limits the activity period of an endothermic Sonoran Desert bee. Functional Ecology 37, 2855–2867. 10.1111/1365-2435.14438

Kamilari, M., Papaioannou, C., Augustinos, A., Spinos, E., Giantsis, I.A., Ramfos, A., Theodorou, J.A., Batargias, C., 2025. From Shell to Sequence: Optimizing DNA Extraction and PCR for Pen Shell Identification. Water 17. 10.3390/w17081162

Kyritsis, G.A., Augustinos, A.A., Livadaras, I., Cáceres, C., Bourtzis, K., Papadopoulos, N.T., 2019. Medfly-Wolbachia symbiosis: genotype x genotype interactions determine host’s life history traits under mass rearing conditions. BMC Biotechnology 19, 96. 10.1186/s12896-019-0586-7

Kyritsis, G.A., Koskinioti, P., Bourtzis, K., Papadopoulos, N.T., 2022. Effect of Wolbachia Infection and Adult Food on the Sexual Signaling of Males of the Mediterranean Fruit Fly Ceratitis capitata. Insects 13, 737. 10.3390/insects13080737

Lee, C.E., 2002. Evolutionary genetics of invasive species. Trends in Ecology & Evolution 17, 386–391. 10.1016/S0169-5347(02)02554-5

Loukeris, T.G., Livadaras, I., Arcà, B., Zabalou, S., Savakis, C., 1995. Gene Transfer into the Medfly, Ceratitis capitata, with a Drosophila hydei Transposable Element. Science 270, 2002–2005.

Mateos, M., Martinez Montoya, H., Lanzavecchia, S.B., Conte, C., Guillén, K., Morán-Aceves, B.M., Toledo, J., Liedo, P., Asimakis, E.D., Doudoumis, V., Kyritsis, G.A., Papadopoulos, N.T., Augustinos, A.A., Segura, D.F., Tsiamis, G., 2020. Wolbachia pipientis Associated With Tephritid Fruit Fly Pests: From Basic Research to Applications. Front. Microbiol. 11. 10.3389/fmicb.2020.01080

McCluney, K.E., 2017. Implications of animal water balance for terrestrial food webs. Current Opinion in Insect Science, Global change biology * Molecular physiology 23, 13–21. 10.1016/j.cois.2017.06.007

McCluney, K.E., Belnap, J., Collins, S.L., González, A.L., Hagen, E.M., Nathaniel Holland, J., Kotler, B.P., Maestre, F.T., Smith, S.D., Wolf, B.O., 2012. Shifting species interactions in terrestrial dryland ecosystems under altered water availability and climate change. Biological Reviews 87, 563–582. 10.1111/j.1469-185X.2011.00209.x

McDermott, E.G., Mullens, B.A., 2014. Desiccation Tolerance in the Eggs of the Primary North American Bluetongue Virus Vector, Culicoides sonorensis (Diptera: Ceratopogonidae), and Implications for Vector Persistence. J Med Entomol 51, 1151–1158. 10.1603/ME14049

Moss, W.E., Crausbay, S.D., Rangwala, I., Wason, J.W., Trauernicht, C., Stevens-Rumann, C.S., Sala, A., Rottler, C.M., Pederson, G.T., Miller, B.W., Magness, D.R., Littell, J.S., Frelich, L.E., Frazier, A.G., Davis, K.T., Coop, J.D., Cartwright, J.M., Booth, R.K., 2024. Drought as an emergent driver of ecological transformation in the twenty-first century. BioScience 74, 524–538. 10.1093/biosci/biae050

O’Neill, S.L., Karr, T.L., 1990. Bidirectional incompatibility between conspecific populations of Drosophila simulans. Nature 348, 178–180. 10.1038/348178a0

Papadogiorgou, G.D., Papadopoulos, A.G., Moraiti, C.A., Verykouki, E., Papadopoulos, N.T., 2024. Latitudinal variation in survival and immature development of Ceratitis capitata populations reared in two key overwintering hosts. Sci Rep 14, 467. 10.1038/s41598-023-50587-2

Sarakatsanou, A., Diamantidis, A.D., Papanastasiou, S.A., Bourtzis, K., Papadopoulos, N.T., 2011. Effects of Wolbachia on fitness of the Mediterranean fruit fly (Diptera: Tephritidae). Journal of Applied Entomology 135, 554–563. 10.1111/j.1439-0418.2011.01610.x

Saridaki, A., Bourtzis, K., 2010. Wolbachia: more than just a bug in insects genitals. Current Opinion in Microbiology 13, 67–72. 10.1016/j.mib.2009.11.005

Schmidt, C.A., Comeau, G., Monaghan, A.J., Williamson, D.J., Ernst, K.C., 2018. Effects of desiccation stress on adult female longevity in Aedes aegypti and Ae. albopictus (Diptera: Culicidae): results of a systematic review and pooled survival analysis. Parasites Vectors 11, 267. 10.1186/s13071-018-2808-6

Shelly, T.E., 2018. Larval host plant influences male body size and mating success in a tephritid fruit fly. Entomologia Experimentalis et Applicata 166, 41–52. 10.1111/eea.12639

Shoukry, A., Hafez, M., 1979. Studies on the Biology of the Mediterranean Fruit Fly Ceratitis Capitata. Entomologia Experimentalis et Applicata 26, 33–39. 10.1111/j.1570-7458.1979.tb02894.x

Simmons, L.W., Lovegrove, M., Du, X. (Bob), Ren, Y., Thomas, M.L., 2023. Humidity stress and its consequences for male pre- and post-copulatory fitness traits in an insect. Ecology and Evolution 13, e10244. 10.1002/ece3.10244

Sinclair, B.J., Saruhashi, S., Terblanche, J.S., 2024. Integrating water balance mechanisms into predictions of insect responses to climate change. Journal of Experimental Biology 227, jeb247167. 10.1242/jeb.247167

StatsCalculators.com - Free Online Statistics Calculators [WWW Document], n.d. URL https://www.statscalculators.com/ (accessed 4.15.26).

Szyniszewska, A.M., Bieszczak, H., Kozyra, K., Papadopoulos, N.T., De Meyer, M., Nowosad, J., Ota, N., Kriticos, D.J., 2024. Evidence that recent climatic changes have expanded the potential geographical range of the Mediterranean fruit fly. Sci Rep 14, 2515. 10.1038/s41598-024-52861-3

Szyniszewska, A.M., Tatem, A.J., 2014. Global Assessment of Seasonal Potential Distribution of Mediterranean Fruit Fly, Ceratitis capitata (Diptera: Tephritidae). PLOS ONE 9, e111582. 10.1371/journal.pone.0111582

Thorat, L., Nath, B.B., 2018. Insects With Survival Kits for Desiccation Tolerance Under Extreme Water Deficits. Front. Physiol. 9. 10.3389/fphys.2018.01843

Wang, Y., Ferveur, J.-F., Moussian, B., 2021. Eco-genetics of desiccation resistance in Drosophila. Biological Reviews 96, 1421–1440. 10.1111/brv.12709

Weldon, C.W., Boardman, L., Marlin, D., Terblanche, J.S., 2016. Physiological mechanisms of dehydration tolerance contribute to the invasion potential of Ceratitis capitata (Wiedemann) (Diptera: Tephritidae) relative to its less widely distributed congeners. Frontiers in Zoology 13, 15. 10.1186/s12983-016-0147-z

Weldon, C.W., Díaz-Fleischer, F., Pérez-Staples, D., 2019. Desiccation Resistance of Tephritid Flies: Recent Research Results and Future Directions, in: Area-Wide Management of Fruit Fly Pests. CRC Press.

Weldon, C.W., Nyamukondiwa, C., Karsten, M., Chown, S.L., Terblanche, J.S., 2018. Geographic variation and plasticity in climate stress resistance among southern African populations of Ceratitis capitata (Wiedemann) (Diptera: Tephritidae). Sci Rep 8, 9849. 10.1038/s41598-018-28259-3

Yen, J.H., Barr, A.R., 1971. New hypothesis of the cause of cytoplasmic incompatibility in Culex pipiens L. [31]. Nature 232, 657–658. 10.1038/232657a0

Zabalou, S., Apostolaki, A., Livadaras, I., Franz, G., Robinson, A.S., Savakis, C., Bourtzis, K., 2009. Incompatible insect technique: incompatible males from a Ceratitis capitata genetic sexing strain. Entomologia Experimentalis et Applicata 132, 232–240. 10.1111/j.1570-7458.2009.00886.x

Zabalou, S., Riegler, M., Theodorakopoulou, M., Stauffer, C., Savakis, C., Bourtzis, K., 2004. Wolbachia-induced cytoplasmic incompatibility as a means for insect pest population control. Proceedings of the National Academy of Sciences 101, 15042–15045. 10.1073/pnas.0403853101

Zacharopoulou, A., Augustinos, A. a., Drosopoulou, E., Tsoumani, K. t., Gariou-Papalexiou, A., Franz, G., Mathiopoulos, K. d., Bourtzis, K., Mavragani-Tsipidou, P., 2017. A review of more than 30 years of cytogenetic studies of Tephritidae in support of sterile insect technique and global trade. Entomologia Experimentalis et Applicata 164, 204–225. 10.1111/eea.12616

Zug, R., Hammerstein, P., 2012. Still a host of hosts for Wolbachia: Analysis of recent data suggests that 40% of terrestrial arthropod species are infected. PLoS ONE 7. 10.1371/journal.pone.0038544

Zwiebel, L.J., Saccone, G., Zacharopoulou, A., Besansky, N.J., Favia, G., Collins, F.H., Louis, C., Kafatos, F.C., 1995. The white gene of Ceratitis capitata: a phenotypic marker for germline transformation. Science 270, 2005–2008. 10.1126/science.270.5244.2005

