## Supplementary material for "Egg-stage desiccation reduces developmental recovery and reveals strain-dependent *Wolbachia*-associated costs in the Mediterranean fruit fly, *Ceratitis capitata*": Suuplementary Files

**Table S1. Full descriptive statistics for baseline control-condition comparisons among the three** *Ceratitis* **capitata strains. D**etailed descriptive statistics for hatching, pupation, and adult emergence under control conditions in the Benakeion, 88.6, and S.10.3 strains, including sample size, mean, standard deviation, median, standard error, minimum, and maximum values for each endpoint.

| Strain | Statistic | Hatching | Pupation on hatched eggs | Pupation on eggs | Emergence on pupae | Emergence on eggs |
| --- | --- | --- | --- | --- | --- | --- |
| Benakeion | Mean | 94.4667 | 97.7333 | 92.2 | 100 | 92.2 |
| Benakeion | SD | 3.86825 | 1.96299 | 1.90526 | 0 | 1.90526 |
| Benakeion | Median | 96.7 | 96.6 | 93.3 | 100 | 93.3 |
| 88.6 | Mean | 73.3667 | 96.2333 | 70.6333 | 98.2333 | 69.4333 |
| 88.6 | SD | 1.823 | 1.30512 | 2.36925 | 3.05996 | 4.44672 |
| 88.6 | Median | 73.7 | 96.1 | 71.9 | 100 | 71.9 |
| S.10.3 | Mean | 58.3333 | 95.7667 | 55.8333 | 93.8667 | 52.5 |
| S.10.3 | SD | 3.81881 | 4.35469 | 3.81881 | 6.89734 | 6.61438 |
| S.10.3 | Median | 57.5 | 96 | 55 | 95.2 | 50 |

**Table S2. Full Dunn pairwise comparison matrix for the effects of egg-desiccation duration on developmental recovery in** *Ceratitis* **capitata.** Adjusted p-values from Dunn’s post hoc pairwise comparisons among the seven egg-desiccation treatments (0, 4, 8, 12, 16, 20, and 24 h) for hatching, pupation on eggs, and adult emergence on eggs, following significant Kruskal–Wallis tests.

| Comparison | Hatching adj. p | Pupation on eggs adj. p | Emergence on eggs adj. p |
| --- | --- | --- | --- |
| 0 vs 12 | 1 | 0.5046 | 0.4408 |
| 0 vs 16 | 0.4117 | 0.0366 | 0.0268 |
| 0 vs 20 | 0.0013* | 0.0003* | 0.0002* |
| 0 vs 24 | 0.0001* | 0* | 0* |
| 0 vs 4 | 1 | 1 | 1 |
| 0 vs 8 | 1 | 1 | 1 |
| 12 vs 16 | 1 | 1 | 1 |
| 12 vs 20 | 0.1795 | 0.7821 | 0.7226 |
| 12 vs 24 | 0.0214 | 0.0599 | 0.0693 |
| 16 vs 20 | 1 | 1 | 1 |
| 20 vs 4 | 0.0007* | 0.0008* | 0.0009* |
| 20 vs 8 | 0.0016 | 0.0018* | 0.0021* |
| 24 vs 4 | 0* | 0* | 0* |
| 24 vs 8 | 0.0001* | 0* | 0.0001* |

**Table S3. Sequential-stage analyses of developmental success after egg-stage desiccation in three** *Ceratitis* **capitata strains.** Kruskal–Wallis and Dunn post hoc results for developmental endpoints expressed relative to the preceding stage, including pupation on hatched eggs and adult emergence on pupae. These analyses assess whether strain-dependent effects persist after successful completion of earlier developmental transitions.

| Endpoint / comparison | Statistic / adj. p | Interpretation |
| --- | --- | --- |
| Kruskal–Wallis: % hatched | χ² = 10.461, p = 0.005* | Significant strain effect. |
| Kruskal–Wallis: % pupation on hatched eggs | χ² = 5.751, p = 0.056 | Not significant. |
| Kruskal–Wallis: % emerged on pupae | χ² = 5.956, p = 0.051 | Not significant. |
| 88.6 vs Benakeion (% hatched) | 0.2292 | Not significant. |
| 88.6 vs S.10.3 (% hatched) | 0.4359 | Not significant. |
| Benakeion vs S.10.3 (% hatched) | 0.0037* | Significant pairwise difference. |

**Table S4. Full logistic regression output for adult survival at 48 h after moderate egg-stage desiccation.** Parameter estimates, standard errors, odds ratios, confidence intervals, and significance values for the binomial logistic regression assessing the effects of strain, egg-desiccation treatment, and their interaction on adult survival at 48 h post-emergence.

| Effect | B / odds ratio | SE / CI | p-value | Comment |
| --- | --- | --- | --- | --- |
| 88.6 vs S.10.3 (48 h survival) | B = 1.279; OR = 3.593 | SE = 0.528; 95% CI 1.276–10.113 | 0.015 | Higher odds of survival than S.10.3. |
| Benakeion vs S.10.3 (48 h survival) | B = 2.197; OR = 9.000 | SE = 0.784; 95% CI 1.937–41.815 | 0.005 | Much higher odds of survival than S.10.3. |
| GLM treatment term | B = 0.113 | 95% CI -0.338 to 0.563 | 0.625 | No treatment effect. |
| GLM interaction term | B = 0.007 | 95% CI -0.179 to 0.193 | 0.944 | No interaction effect supported by the current output. |

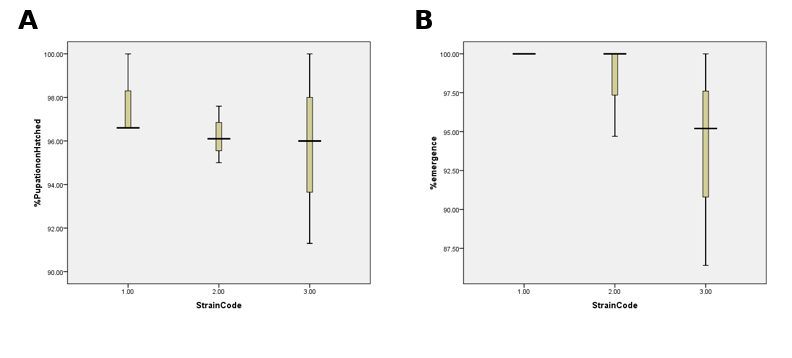
**Figure S1. Conditional-stage developmental performance of the three** *Ceratitis* **capitata strains under control conditions.** Pupation expressed relative to the number of hatched eggs and adult emergence expressed relative to the number of pupae for the Benakeion, 88.6, and S.10.3 strains under non-desiccating conditions. In contrast to the clear differences observed for egg-based endpoints in the main text, conditional-stage survival is comparatively similar among strains, indicating that the principal baseline divergence is concentrated at the egg-hatching stage.

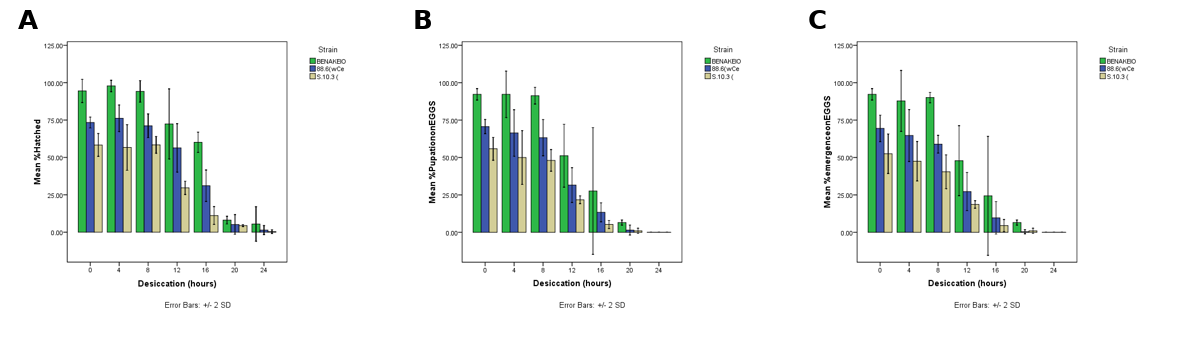

**Figure S2. Effects of egg-stage desiccation duration on developmental recovery in three** *Ceratitis* **capitata strains.** Hatching, pupation, and adult emergence calculated relative to the initial number of eggs after egg desiccation for 0, 4, 8, 12, 16, 20, or 24 h. Across all three strains, developmental performance remains comparatively stable during short desiccation exposures and declines progressively with increasing duration, with the strongest loss of developmental recovery observed after prolonged exposure, particularly at 20–24 h. Benakeion consistently shows the highest recovery, S.10.3 the lowest, and 88.6 intermediate performance.

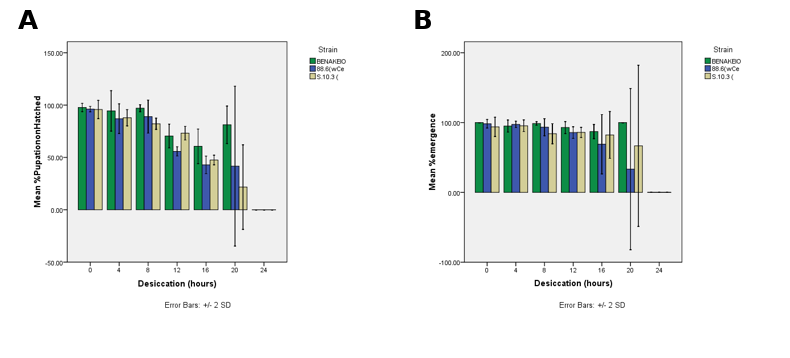

**Figure S3. Conditional-stage developmental success after egg-stage desiccation in three** *Ceratitis* **capitata strains.** Pupation expressed relative to hatched eggs and adult emergence expressed relative to pupae following egg desiccation for 0, 4, 8, 12, 16, 20, or 24 h. These conditional-stage endpoints show that once individuals successfully complete hatching, later developmental transitions are less strongly differentiated among strains than egg-based recovery, supporting the interpretation that strain-dependent desiccation effects are expressed primarily at the embryonic and hatching stages.
